# Multi-omic characterization of antibody-producing CHO cell lines elucidates metabolic reprogramming and nutrient uptake bottlenecks

**DOI:** 10.1101/2023.09.13.557626

**Authors:** Saratram Gopalakrishnan, William Johnson, Miguel A. Valderrama-Gomez, Elcin Icten, Jasmine Tat, Fides Lay, Jonathan Diep, Natalia Gomez, Jennitte Stevens, Fabrice Schlegel, Pablo Rolandi, Cleo Kontoravdi, Nathan Lewis

## Abstract

Characterizing the phenotypic diversity and metabolic capabilities of industrially relevant manufacturing cell lines is critical to bioprocess optimization and cell line development. Metabolic capabilities of the production hosts limit nutrient and resource channeling into desired cellular processes and can have a profound impact on productivity but cannot be directly inferred from measured data such as spent media concentrations or transcriptomics. Here, we present an integrated multi-omic characterization approach combining exo-metabolomics, transcriptomics, and genome-scale metabolic network analysis and apply it to three antibody-producing Chinese Hamster Ovary cell lines to reprogramming features associated with high-producer clones and metabolic bottlenecks limiting product production in an industrial bioprocess. Analysis of individual datatypes revealed a decreased nitrogenous byproduct secretion in high-producing clones and the topological changes in peripheral metabolic pathway expression associated with phase shifts. An integrated omics analysis in the context of the genome-scale metabolic model elucidated the differences in central metabolism and identified amino acid utilization bottlenecks limiting cell growth and antibody production that were not evident from exo-metabolomics or transcriptomics alone. Thus, we demonstrate the utility of a multi-omics characterization in providing an in-depth understanding of cellular metabolism, which is critical to efforts in cell engineering and bioprocess optimization.

## 1. Introduction

Cell line engineering and process optimization are two major challenges faced by the bioprocessing industry in the quest to achieve sustainable production of high-quality products (Tihanyi and Nyitray, 2020). Because the development of engineered strains capable of achieving high product titers is time consuming and costly (Nielsen and Keasling, 2016), there is an imminent need to develop intelligent approaches to cell and process engineering to accommodate existing and emerging products. Systems biology offers a powerful toolkit to characterize cellular metabolism, reveal traits associated with high-producing strains, and elucidate cellular responses to bioreactor conditions (Campbell et al., 2017). Transcriptomic analyses have been applied to uncover the mechanisms governing protein synthesis and secretion in yeast (Huang et al., 2015), and identify overexpression targets associated with cell growth, glycosylation, protein production, and anti-apoptotic factors in CHO cells (Fischer et al., 2015). On the other hand, metabolomic data has been used to reveal the existence of multiple distinct cellular states over the course of the bioprocess with an exponential phase and a stationary phase being the primary phases in which cells prioritize growth and antibody production, respectively (Ahn and Antoniewicz, 2011; Dean and Reddy, 2013; Templeton et al., 2013). Sub-phases can involve lactate consumption (Mulukutla et al., 2012), increases in cell size (Pan et al., 2017), and declining viability (Templeton et al., 2013). Despite these advances, protein yields using CHO cells remain far below the maximum possible theoretical specific productivity (less than 57%) despite recent advancements in cell line development (Hefzi et al., 2016). Because individual data types (e.g., metabolomics from spent media, transcriptomics, or fluxomics) provide a partial picture of dynamic cellular behavior in the bioreactor (Haas et al., 2017), efforts geared towards improving specific attributes such as cell growth, cell viability, or antibody productivity focus on improving the performance of only a subset of the phases in a bioprocess. A multi-omics study that encompasses exo-metabolomics, transcriptomics and fluxomics will provide a holistic understanding of cellular metabolism in a bioreactor by elucidating nutrient uptake, pathway utilization, and metabolic priorities of the cell in the various phases and reveal specific engineering targets to improve the antibody titer.

Here, we present a multi-omics analysis workflow to demonstrate the phenotypic and metabolic diversity of three producing CHO cell lines (including a non-clonal pools and a high-producing clone derived from each) and identify metabolic bottlenecks that cannot be identified using any one type of data alone. First, we analyze the overall nutrient consumption and product yield in the bioreactor and show that high-producing clones do not necessarily need to grow to high cell densities, and that low cell density clones can achieve high titers through high specific antibody productivity. We built cell line-specific metabolic models with gene expression data using a published extraction workflow (Gopalakrishnan et al., 2023) using the mCADRE algorithm (Wang et al., 2012) and StanDep (Joshi et al., 2020) for expression thresholding and observed differences in model content involving lipid metabolism, glycosylation, and intracellular transport pathways. Since energy- and precursor-producing pathways (e.g., central metabolism, amino acid catabolism, and nucleotide biosynthesis) were conserved across all cell lines in all phases, we probed intracellular flux distributions using Monte-Carlo flux sampling after overlaying the uptake and secretion rates inferred from exometabolomics data on to the constructed phase- and cell line-specific models and found that glycolysis was primarily driven by glucose uptake and the TCA cycle was driven by glutaminolysis and degradation of acetogenic amino acids such as lysine and valine. From a model-based analysis, we highlight which amino acids become limiting in each process phase and describe the observed rewiring of nitrogen metabolism that enabled efficient amino acid usage over the course of the bioprocess in high-producing cell lines.

## 2. Methods

### 2.1 Cell culture, exo-metabolomic, and protein concentration quantification

Three high-expressing clonally derived Chinese hamster ovary (CHO) cell lines (Clone D, E, G) and their corresponding progenitor non-clonal pools (Pool D, E, G, respectively), each expressing a non-glycosylated antibody were used in this study. Cells were thawed and expanded to inoculate fed-batch spin tube bioreactors (TPP, Trasadingen, Switzerland) containing proprietary chemically defined growth media in triplicate at target viable cell density (VCD) of 8×10^5^ cells/mL and working volume of 30 mL. Cells were incubated at 36°C, 5% CO2, 85% relative humidity and shaken at 225 rpm with 50 mm orbital diameter in a large-capacity ISF4-X incubator (Kuhner AG, Basel, Switzerland). Cultures were fed a single bolus feed of proprietary feed media on days 3, 6, and 8. VCD (measured using Vi-Cell BLU cell counter, Beckman Coulter, Brea, CA), spent-media exo-metabolites (measured using BioProfile FLEX2, Nova Biomedical, Waltham, MA), spent-media amino acids (measured with Ultra Performance liquid chromatography analysis, UV detection and AccQ-Tag Derivatization Chemistry, Waters, Milford, MA) and titer (measured using Protein A high performance liquid chromatography analysis, Protein A, Waters, Milford, MA) were analyzed throughout the culture duration of ten days.

### 2.2 Transcriptomic quantification and data processing

Cells from clones D, E, and G were collected and cryo-preserved on Day 0 (the culture prior to inoculation) and Day 7 of the fed-batch process for transcriptomic analysis. 1×10^6^ to 1×10^7^ cells per culture were spun-down and resuspended in proprietary culture media with 10% DMSO (Sigma-Aldrich, St. Louis, MO) and frozen at −80 °C in cryovials (Thermo Scientific, Waltham, MA). Cryovials were thawed at 36°C and total RNA was extracted (RNeasy Micro Kit, Qiagen, Hilden, Germany) in one batch. cDNA libraries were generated NEBNext Ultra RNA Library Prep for Illumina following manufacturer’s instructions (NEB, Ipswich, MA, USA) and sequenced on a HiSeq 4000 using a 2×150bp Paired End configuration (Illumina, La Jolla, CA), targeting 50 million paired-end reads per library. Sequence reads were trimmed to remove adapter sequences and nucleotides with poor quality using Trimmomatic v.0.36 (Bolger et al., 2014). The trimmed reads were mapped to the Chinese Hamster PICR reference genome (Rupp et al., 2018) using the STAR aligner v.2.5.2b (Dobin et al., 2013). Unique gene hit counts were calculated by using featureCounts from the Subread package v.1.5.2 (Liao et al., 2014). Only unique reads that fell within exon regions were counted.

### 2.3 Concentration data processing and flux calculations

The growth rate, specific antibody productivity, and specific uptake and secretion rates for all quantified metabolites were computed as described earlier (Templeton et al., 2013). The process was divided into four phases based on feeding intervals. Phase 1 was between days 0 – 3, phase 2 was between days 3 – 6, phase 3 was between days 6 – 8, and phase 4 was between days 8 – 10. The growth rate and the specific uptake and secretion rates in phase *k* were computed by fitting the concentration data to the following set of ODEs:

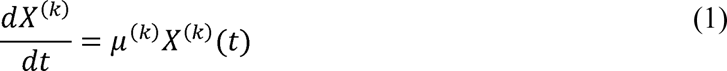

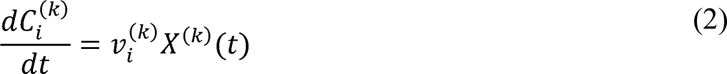

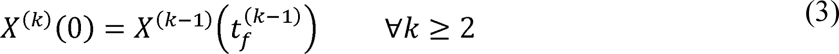

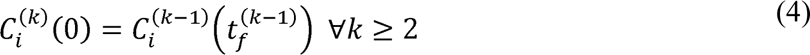

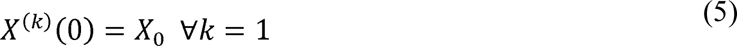

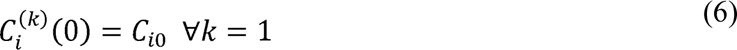

In the above equations, 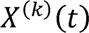 is the cell density during phase 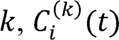 is the concentration of metabolite *i* during phase 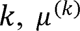 is the growth rate during phase *k*, and 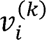 is the specific uptake rate/productivity of metabolite *i* during phase *k*. We solve a least-squares regression problem to obtain 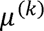 and 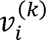 for all metabolites in all phases. The computed fluxes are reported in Supplementary Table ST2.

### 2.4 Generating context-specific models for producer clones

Context-specific models for the pool and clone cultures for each phase were extracted using the workflow described earlier (Gopalakrishnan et al., 2023). First, the measured uptake and secretion rates were imposed as bounds in the *i*CHO1766 genome-scale metabolic model (Hefzi et al., 2016) and the inactive reactions were identified using flux variability analysis (FVA) (Mahadevan and Schilling, 2003). Transcriptomics data integration was performed using the mCADRE algorithm (Wang et al., 2012). Thresholds generated using StanDep (Joshi et al., 2020) were applied to the gene expression data, from which, ubiquity scores were calculated. A cutoff of 1 was used to classify reactions into core and non-core reactions. All metabolites with available extracellular measurements were included in the set of core reactions. Since mCADRE shows the least variability in model content (Gopalakrishnan et al., 2023), ensembles of 20 models were generated in each case and then combined to account for potential alternate solutions. Model similarity was quantified using the Jaccard Index defined as follows:

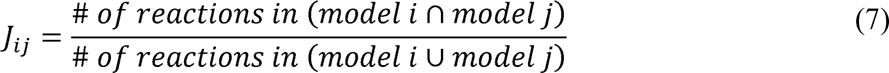

### 2.5 Monte-Carlo flux sampling

Intracellular flux distributions for the extracted models were obtained using the Coordinate Hit-and-Run Monte-Carlo flux sampling approach (Haraldsdóttir et al., 2017). Because metabolic models inherently contain thermodynamically infeasible cycles (TICs) (Schellenberger et al., 2011), reactions associated with these cycles must be inactivated to ensure that obtained flux distributions do not violate the second law of thermodynamics. A hallmark of such cycles is that they afford flux distributions that are several orders of magnitude higher than the uptake and secretion rates of metabolites in the model. We circumvent this issue by (i) identifying the most relevant set of fluxes to be sampled and (ii) minimizing the flux through all reactions associated with TICs. First, we decompose all fluxes in the metabolic models into “free” and “dependent fluxes” (Famili and Palsson, 2003; Wiechert et al., 1997) and express the dependent fluxes in terms of the free fluxes using the null space matrix. This decomposition reveals that every flux is a combination of an extracellular uptake/secretion flux, intracellular fluxes that are a part of alternate routes, and intracellular fluxes that may be involved in TICs (Chan et al., 2018). We then set all extracellular uptake/secretion rates to zero in the model and perform FVA to identify all free fluxes with non-zero bounds. These reactions are potentially involved in TICs because they allow flux in the absence of actual metabolite flows within the metabolic model. We then use parsimonious flux balance analysis (pFBA) (Lewis et al., 2010) to selectively minimize the flux through free fluxes associated with TICs. Free fluxes that carry no flux in the pFBA solution are inactivated in the context-specific metabolic model. This results in a smaller subset of free fluxes that can be meaningfully sampled using Monte-Carlo flux sampling. We generated 10,000 flux samples per condition as described by Haraldsdóttir et al. (2017) with the pFBA solution as the starting point and 100 steps per coordinate. The mean flux and the standard deviation for each reaction are reported in Supplementary Table ST4.

## 3. Results

### 3.1. A multi-omics workflow to characterize cellular metabolism in a bioreactor

Diverse types of biological data and analysis approaches can be used for characterizing the cells and bioprocesses in biomanufacturing (Figure 1). Exo-metabolomics (time-course metabolite concentration data measured from spent media) is the most widely used data type in the bioprocessing industry. Statistical analysis of exo-metabolomics data reveals overall nutrient consumption and product formation in the bioprocess and provides valuable insights into process yield, nutrient depletion, and correlations between nutrient consumption and product yields to guide media design. However, this approach may miss the dynamics within the bioreactor arising from cell state and metabolic shifts that affect culture performance and process productivity.

**Figure 1:**
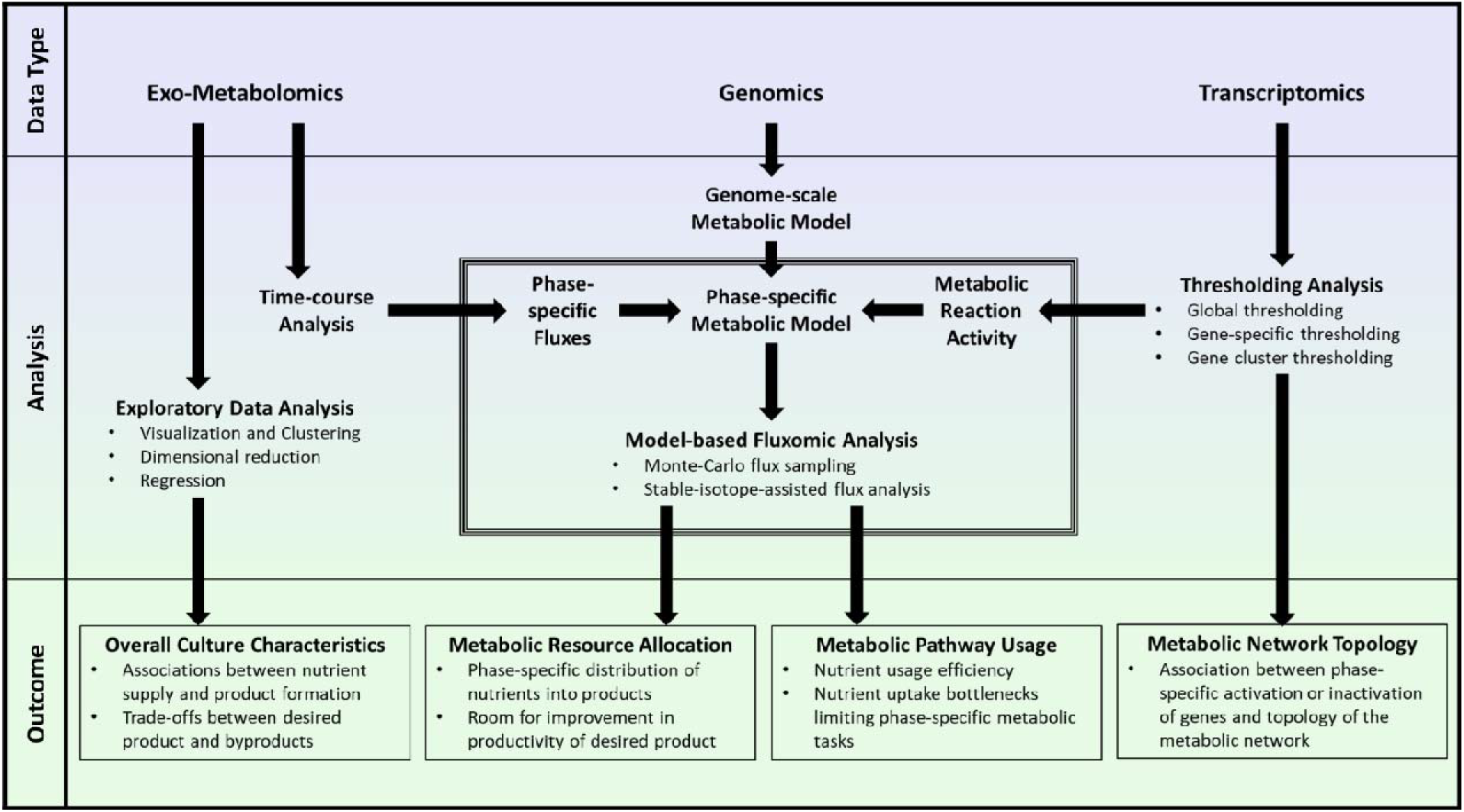
Multi-omics analysis workflow showing the various data types and analysis methods to characterize cellular metabolism in a bioreactor.

Transcriptomics can provide a means to study phase shifts by quantifying changes in gene expression in different process phases, which can further elucidate the metabolic network topology using several available algorithms (Opdam et al., 2017; Robaina Estevez and Nikoloski, 2014), explore phase-specific metabolic tasks (Richelle et al., 2019), and identify engineering targets to improve cell viability in the bioprocess (Fischer et al., 2015). In contrast, fluxomics using either flux sampling (Fallahi et al., 2020) or stable-isotope-assisted metabolic flux analysis (Sacco and Young, 2021), requires a metabolic model derived from the genomic sequence (Mendoza et al., 2019) and can quantify intracellular pathway usage and reveal process bottlenecks. However, such approaches are computationally prohibitive with comprehensive metabolic models. Constraining metabolic models using uptake and secretion rates (inferred from exo-metabolomics data) and reaction inactivity (inferred from transcriptomics data) greatly reduced the size of genome-scale metabolic models by as much as 79% in this study, thereby permitting the use of existing flux elucidation methods.

Here we deployed a workflow that integrates these three datasets, leveraging the strengths of each data type, as well as the synergy between the data types. Through this, we obtain an in-depth understanding of cell behavior over the course of the bioprocess (Figure 1). We use this workflow to quantify the biological differences between three Chinese Hamster Ovary (CHO) cell lines (cell lines D, E, and G), each consisting of one high-producing clonally derived cell line (hereafter referred to as clone-D, clone-E, and clone-G) and corresponding progenitor non-clonal stable pools (hereafter referred to as pool-D, pool-E, and pool-G) with respect to nutrient consumption, metabolic state shifts, transcriptomic, and fluxomic changes over the course of the bioprocess. Taken together, these changes reveal the key phenotypic features of several high-producing clones and the amino acid uptake bottlenecks affecting cell growth and antibody production in a fed-batch bioreactor.

### 3.2. Exometabolomic analysis reveals phenotypic diversity among high-producing clones

We analyzed exo-metabolomic data (metabolite concentration profiles over the course of the bioprocess) to provide insights into the phenotype of high-producing clones and their potential impacts on overall yield and productivity of the bioprocess. To this end, we first computed the total biomass formation, nutrient consumption, and resource allocation of clones, compared to the corresponding pool. No consistent trend was observed between final cell density and total antibody production across all three clones and respective parental pools (Figure 2A). Consistent with previously published features of high-producer clones (Templeton et al., 2013), clone-G showed an increased final cell density compared to its parental pool. Meanwhile, clone-D showed no significant difference in final cell density. Interestingly, clone-E had a lower final cell density compared to its parental pool.

**Figure 2A:**
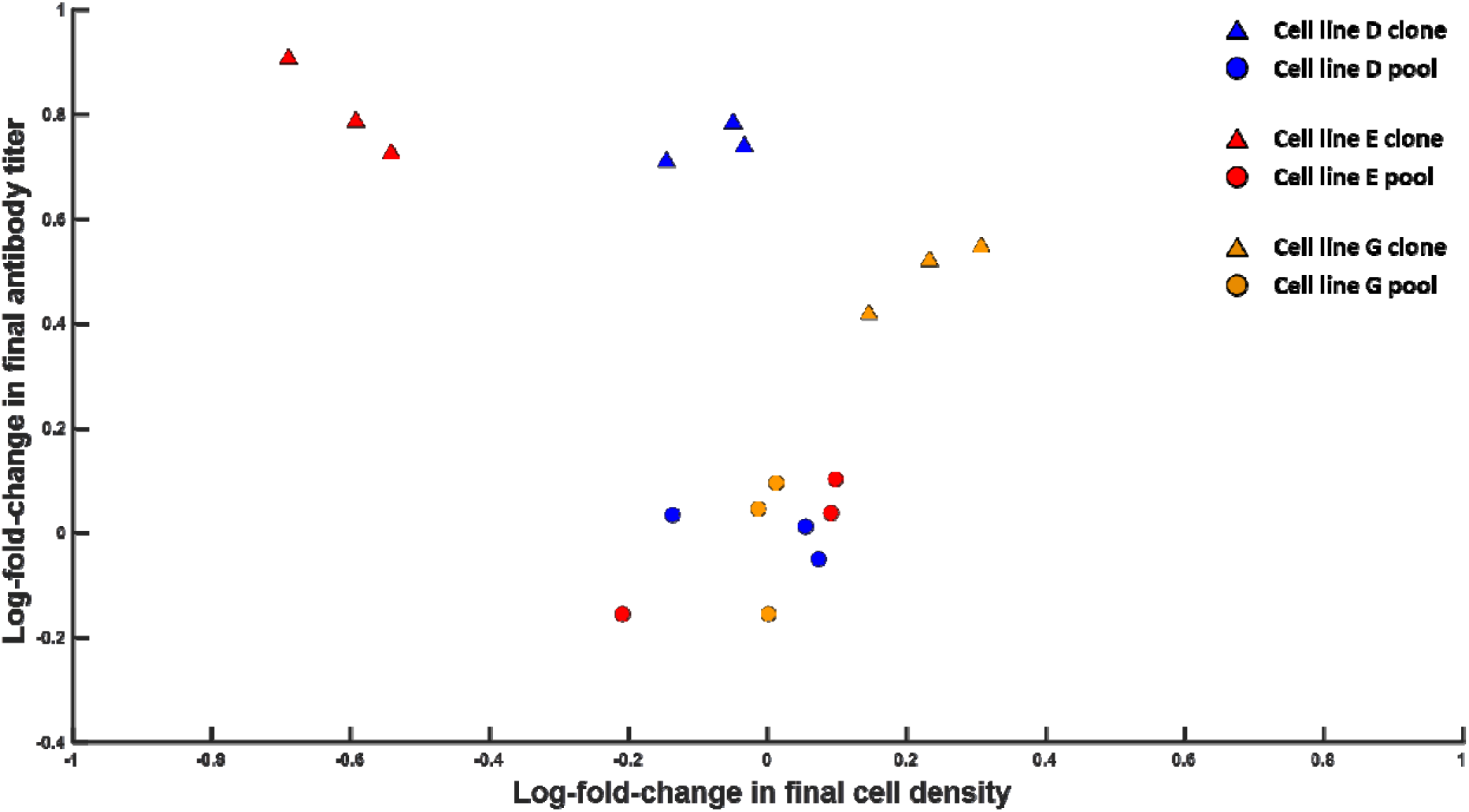
Process characteristics of pool and clone cultures in three producer cell lines. The x-axis represents the log-fold-change in the final cell density of various clones relative to their parental pools. The y-axis represents the log-fold-change in the final antibody titer in producer clones compared to their respective pool cultures. Experimental data was measured in triplicates. All pool culture data is clustered around the origin (0,0), which represents no change in final cell density and antibody titer relative to the parental pools for each cell line.

Given the differences in cell density and productivity, we investigated how nutrient consumption changes between the clones and pools. Thus, we performed a partial least squares (PLS) analysis using the total antibody produced in each culture as the response (Y) and total change in all other quantities (cell density, glucose, lactate, ammonia, and 20 amino acids) as the predictors (X). PLS regression revealed how changes in nutrient uptake and product secretion correlate with overall antibody production. These correlations indicate potential metabolic couplings or trade-offs between antibody and other products in a bioprocess (biomass, lactate, ammonia, and secreted amino acids) (Figure 2B). Glucose consumption was not significantly different between pool and clone cultures, but instead accounted for internal variation within the pool and clone cultures. Lactate production correlated positively with final cell density (a measure of total biomass production) in all three cell lines, with clone-G producing 33% more lactate, and clone-E producing 15% less lactate than the pool-G and pool-E, respectively.

**Figure 2B:**
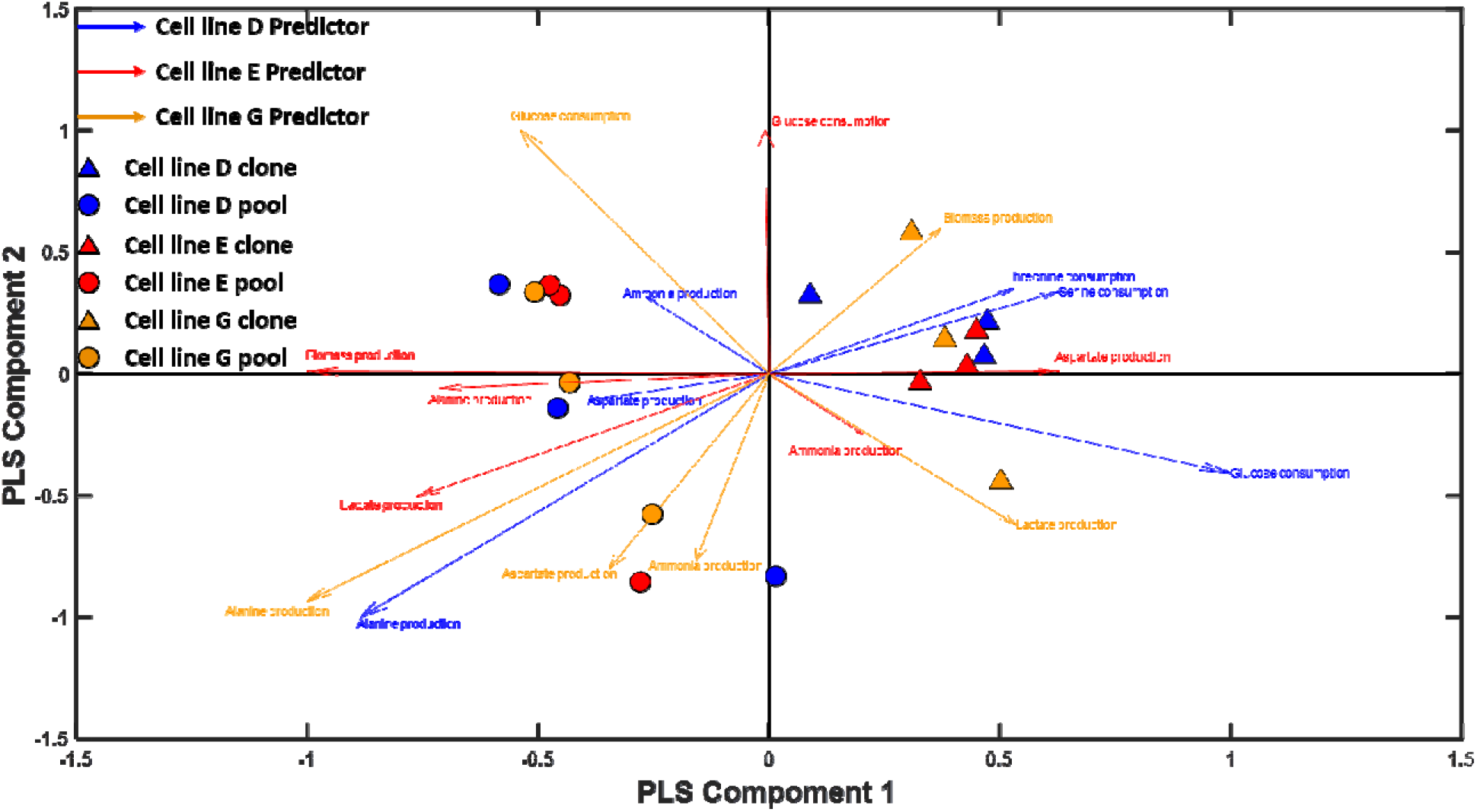
Partial Least Squares (PLS) regression analysis of consumed nutrients and secreted products in cell lines D, E, and G. The markers show the data points projected onto the first two PLS components and the arrows show the contribution of various factors in the first two PLS components.

Changes in the consumption of serine, threonine and production of ammonia, alanine, and aspartate strongly correlated with antibody production (final process titer in the fed-batch process) in the three cell lines (stable pools and clones). Alanine production was consistently lower in the clones compared to the pools for all cell lines (19% lower in clone-D, 55% lower in clone-E, and 39% lower in clone-G). The influence of the other identified factors was largely cell line specific. Antibody production was positively correlated with cell growth in cell line G, highlighting a limited competition between these two metabolic tasks. Cell line E, however, shows a trade-off between more growth in the pool cultures and increased antibody production in the clones. Cell lines D and G showed lower aspartate and ammonia production and higher valine, threonine, and serine consumption in the clone cultures compared to the pool cultures (Supplementary Table ST1). Thus, a rewiring of nitrogen metabolism more efficiently channeled nitrogen into desired products, a trait associated with high-producer clones in this study.

### 3.3. Transcriptomics-based metabolic models elucidate phase- and cell line-specific network topologies

To understand the metabolic state shifts underpinning the reprogramming of nitrogen utilization in clones, we first looked at changes in gene expression over the course of the process and its impact on metabolism. We divided the fed-batch process into four phases based on the feeding schedule (see Methods section 2.3), i.e., the Early Exponential Phase (EXP) between day 0 and day 3, the Lactate Co-utilization Phase (LCP) from day 3 to day 6, the Antibody Production Phase (ABP) between days 6 and 8, and the Late Stationary Phase (LSP) from day 8 to day 10. We built context-specific models for each case using RNA-Seq data (see Methods) and iCHO1766 as the base genome-scale metabolic model to identify and retain active reactions. We evaluated the similarity of the models using the Jaccard Index of included reactions and found that models clustered primarily by process phase (Figure 3A). Within each phase, models corresponding to cell lines D and G were more similar than models for cell line E. Differences in uptake and secretion rates between pool and clone cultures for any cell line did not influence reaction content in phase-specific cell line models. 1,598 reactions were conserved across all models and the differences stemmed from 1,133 reactions that account for 20.5% of the model content.

**Figure 3A:**
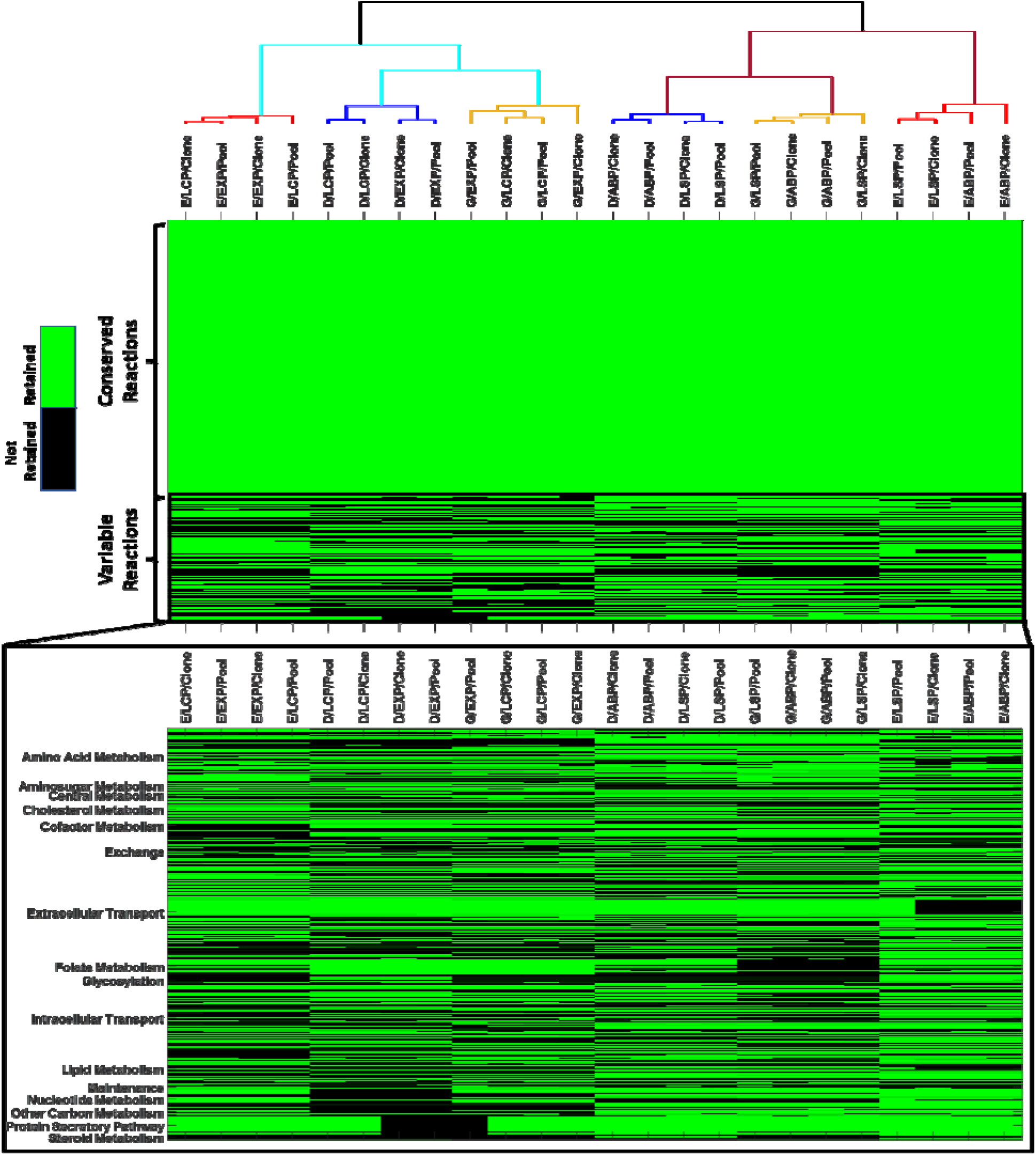
Comparison of reaction content between models extracted for pool and clone cultures of cell lines D, E, and G in all four phases. The heat map shows whether a specific reaction from each pathway is retained in any given model. Green color indicates that the reaction is retained in the model and black color indicates that the reaction is removed during model extraction. In the dendrogram, cell line D is represented in blue, E in red, and G in orange. The cyan lines correspond to the growth phase models (early exponential and lactate co-utilization phases) whereas the maroon lines correspond to the non-growth phases (antibody production and the late stationary phases).

A pathway-level inspection of phase-specific models revealed the antibody production pathway, folate metabolism, and extracellular transport reactions to be primary differences in model content (Figure 3A). The early exponential and lactate co-utilization phase models for cell lines D and G did not contain the antibody production pathway because it was inactivated based on titer measurements for those phases. However, all models retained the pathways necessary to synthesize biomass precursors (nucleotides, triglycerides, cofactors, and cholesterol) to support growth. The nucleotide biosynthesis pathway contained 161 conserved reactions of which 107 were mCADRE-defined core reactions and 54 reactions were coupled to core reactions. An additional 40 reactions were conditionally coupled to core reactions in some models. Lipid metabolism is the largest pathway in iCHO1766 (1,211 reactions), but 845 reactions were blocked or associated with poorly expressed genes. Of the remaining 366 reactions, 302 reactions were in all models because they were coupled to the 84 core reactions and were required to synthesize triacylglycerol and other phospholipids (phosphatidylinositol, phosphatidylserine, phosphatidylethanolamine, and phosphatidylcholine). Hyaluronan metabolism (5 reactions) and keratan sulfate (120 reactions) were fully retained in all models. Although only 20 reactions from keratan sulfate metabolism were core reactions, the remaining reactions were retained because they were coupled to the core reactions. Central metabolism was highly conserved and highly expressed in all extracted models. For amino acid metabolism, 261 reactions (of 574 reactions in iCHO1766) were supported by gene expression data. Of these, 46 reactions were core reactions (associated with highly expressed genes), suggesting that amino acid degradation pathways were primarily retained to metabolize the essential amino acids in excess of that required for cell growth and antibody production. Overall, these findings highlight the differences in peripheral metabolism arising from differences in gene expression, but do not provide any insights into the activity of conserved pathways from central and amino acid metabolism. This motivates the need for flux sampling to investigate the metabolic differences across different conditions.

**Figure 3B:**
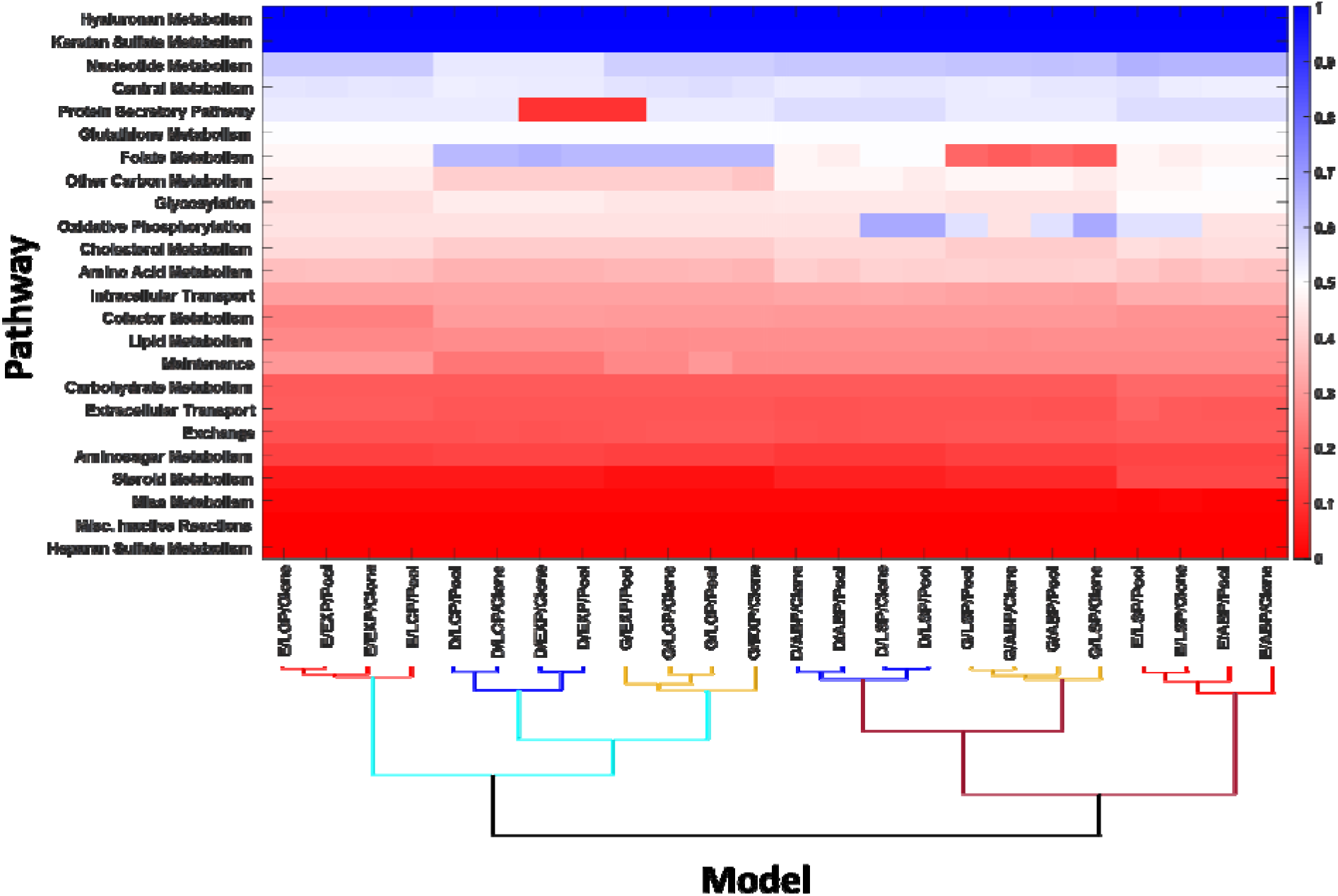
Fractional retention of pathways in extracted context-specific models. A value of 1 indicates that all reactions from the pathway are retained in the context-specific model and a value of 0 indicates that the pathway is completely removed. In the dendrogram, cell line D i represented in blue, E in red, and G in orange. The cyan lines correspond to the growth phase models (early exponential and lactate co-utilization phases) whereas the maroon lines correspond to the non-growth phases (antibody production and the late stationary phases).

### 3.4. Switching from cell growth to antibody production is accompanied by a metabolic shift

While a transcriptomic analysis reveals differences in metabolic network topology across clones and phases, it does not quantify the magnitude of flux, direction of flows, and split ratios within key energy- and precursor-producing pathways within central metabolism. A better understanding of cellular metabolism can be obtained by simulating metabolic fluxes using Monte-Carlo flux sampling on the various constructed context-specific models. A prerequisite for flux sampling is the availability of uptake and secretion rates for all measured metabolites. Analysis of uptake and secretion rates in each process phase revealed significant features of each phase. The EXP is marked by the highest growth rates, glucose uptake, and amino acid consumption in all cultures. The LCP shows a shift from lactate production to lactate consumption. The ABP shows the peak antibody specific productivity in all cultures, whereas the LSP shows an overall decline in metabolic activity marked by a reduction in nutrient uptake and antibody secretion as well as stagnated cell growth. Principal Component Analysis of the computed fluxes showed that the first two principal components explained 94% of the observed variance and that scores are primarily separated based on process phase (Figure 4A). Meanwhile, the transition from the early exponential phase to the lactate co-utilization phase is seen in principal component 2, which entails a reduction in growth rate, a reduction in glucose uptake, a reduction in glycine secretion, and a shift from lactate secretion to lactate consumption. The transition from the lactate co-utilization phase to the antibody production phase was associated with an increase in antibody productivity, which is the main factor within the first principal component (Supplementary Figure S1). Most cultures reverted to lactate production in this phase. Overall, this analysis reveals the phenotypic changes associated with a shift from biomass production to antibody production upon phase shift in the bioprocess.

**Figure 4A:**
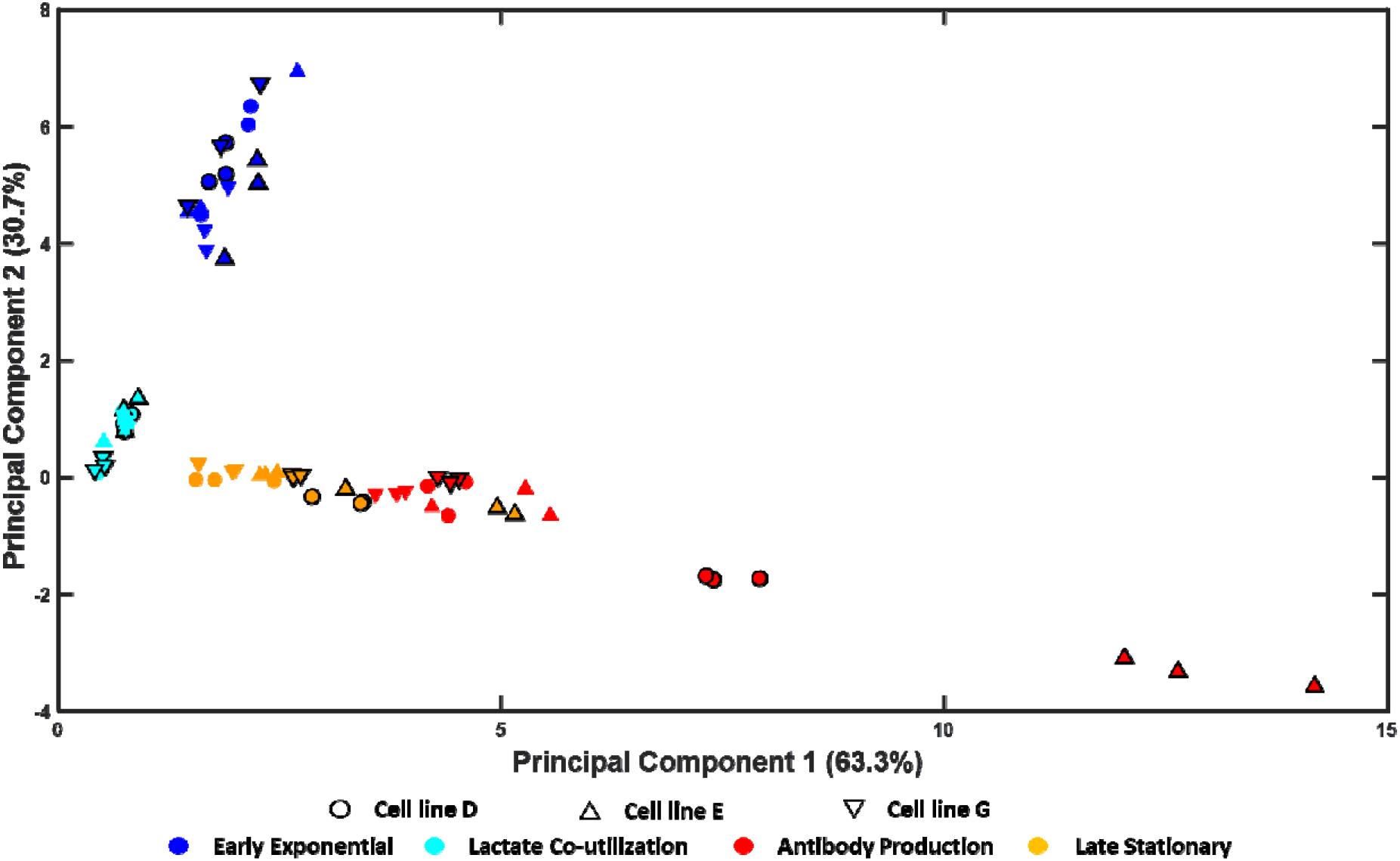
Principal Component Analysis plot showing changes in metabolism over the course of the fed-batch culture. The markers with bolded black edges correspond to the clone cultures whereas the markers without a black edge correspond to the pool cultures. The early exponential and antibody production phases show the largest intra-group variance due to differences in glucose uptake in the early exponential phase and substantial differences in clonal antibody specific productivity in the antibody production phase.

To better understand the differences in clonal pathway usage and explain the overall process changes inferred using exo-metabolomics data, we constrained the previously generated phase-specific metabolic models with the computed uptake and secreted rates and performed Monte-Carlo flux sampling to identify the most likely flux distributions afforded by the metabolic model. We visualized the computed flux distributions using nonmetric multidimensional scaling (Figure 4B) and found that the samples clustered into three distinct groups: the early exponential phase, the lactate co-utilization phase, and a single cluster containing the antibody production and late stationary phases. Unlike with PCA of uptake and secretion rates, the large intra-group variance within the EXP arose from differences in metabolism between pool and clone pathway usage as well as from diversity among high-producing clones, primarily from central metabolism (Figure 5A). Pool-D and pool-G metabolized glucose via the EMP pathway (referred to as UPPER_GLYC in Figure 5). As a result of this, the non-oxidative pentose phosphate pathway (referred to as NOXPPP in Figure 5) operated from triose phosphates (G3P) to pentose phosphates (R5P) to meet the biosynthetic demands for nucleotide biosynthesis. While clone-D was similar to these two pools, clone-G shunted 17.6% of the consumed glucose into the pentose phosphate pathway (referred to as OXPPP in Figure 5), thereby meeting the biosynthetic demands for nucleotide biosynthesis and reversing the direction of flux through NOXPPP to metabolize the excess R5P. In contrast, clone-E showed the reverse trend by diverting the flux through OXPPP in pool-E through the EMP pathway. Lactate yield from glucose increased by 10% in clone-D and by 114% in clone-G compared to pool-D and pool-G, respectively. In clone-G, the anaplerotic pathways (PEPCK and ME) channeled flux from the TCA cycle into lower glycolysis, allowing lactate yield to exceed the theoretical maximum of 2 mol/mol-glucose from glycolysis alone. On the other hand, lactate production in clone-E was lower and the model predicted that flux was instead channeled into glycogen and glycan precursor synthesis. The TCA cycle was fueled by the catabolism of glutamine and asparagine with reductive carboxylation generating the necessary Acetyl-CoA for fatty acid biosynthesis.

**Figure 4B:**
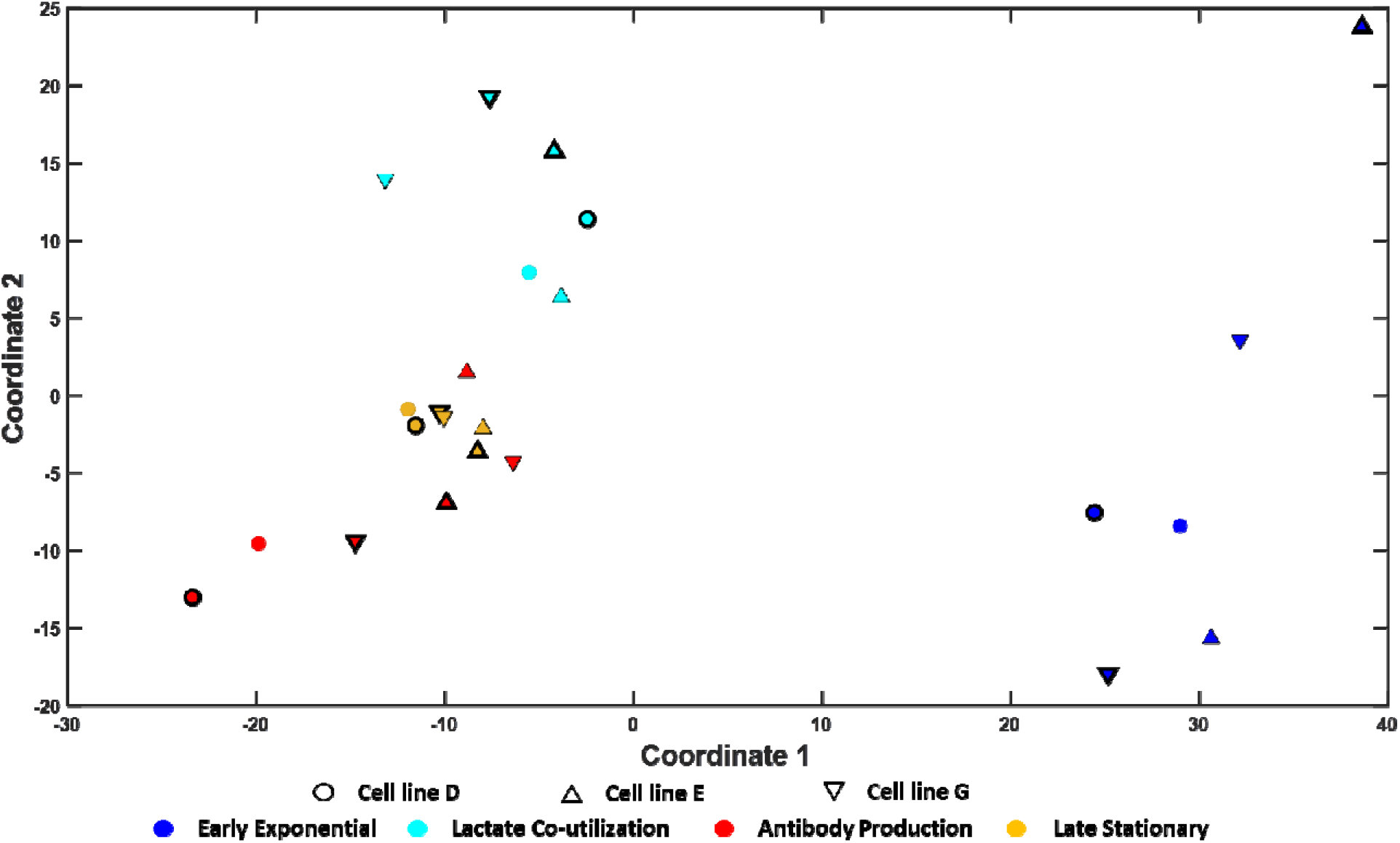
Visualizing intracellular distributions using nonmetric multidimensional scaling. The data separates out into three distinct clusters representing the early exponential phase, the lactat co-utilization phase, and a single cluster containing the antibody production and late stationary phases. Intra-cluster variance represents the differences in metabolism between pool and clones across all cell lines.

**Figure 5:**
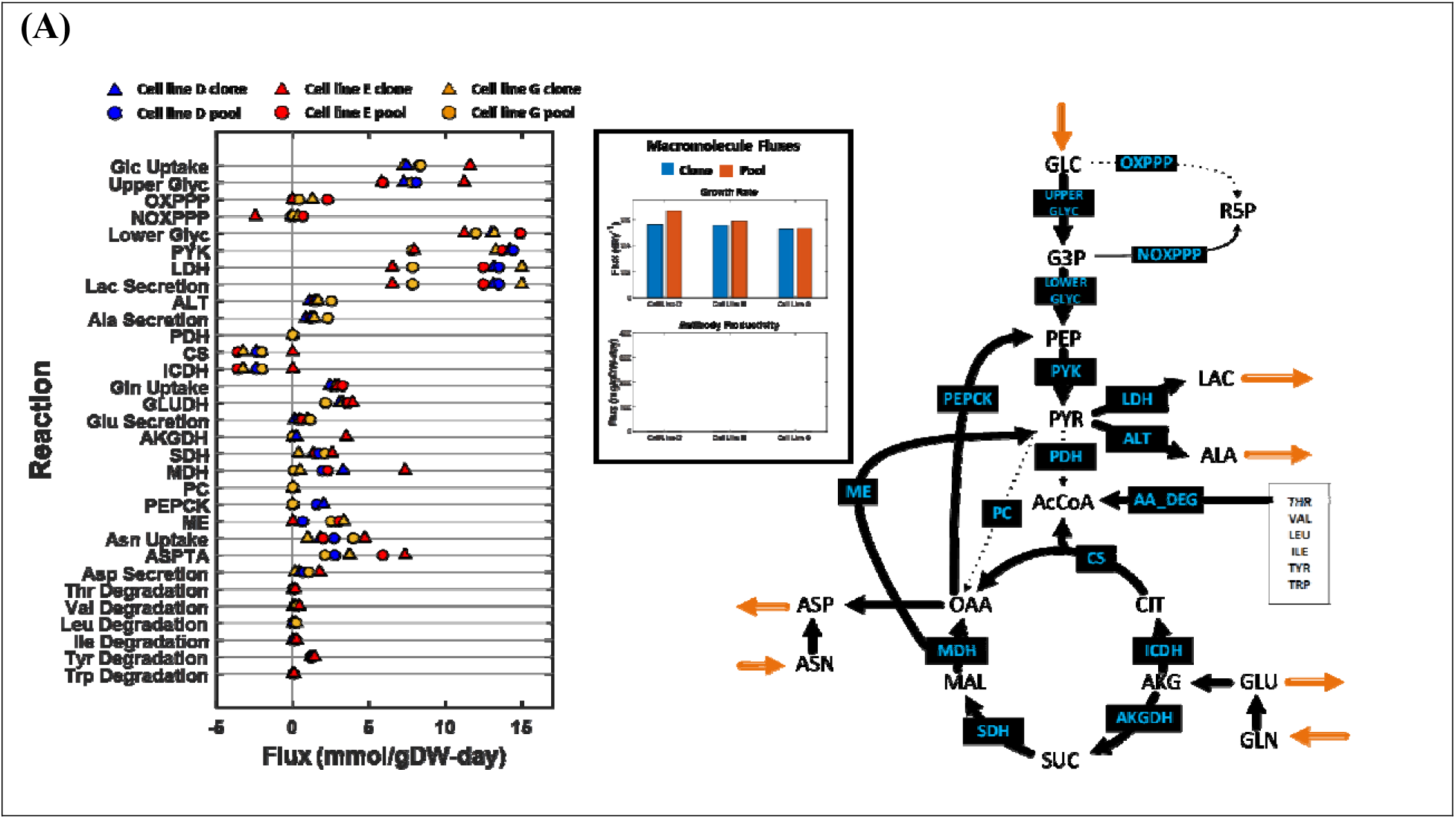
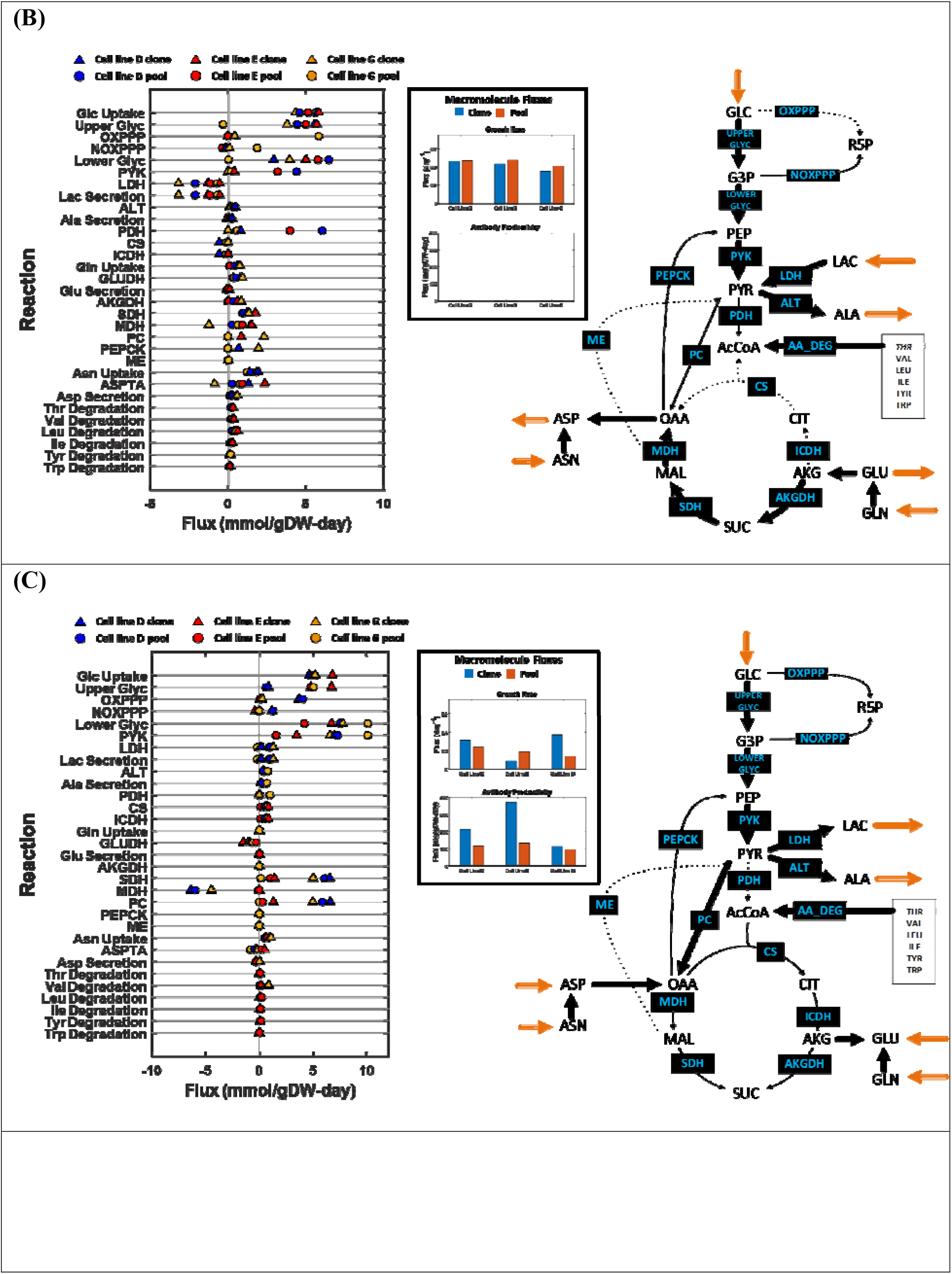
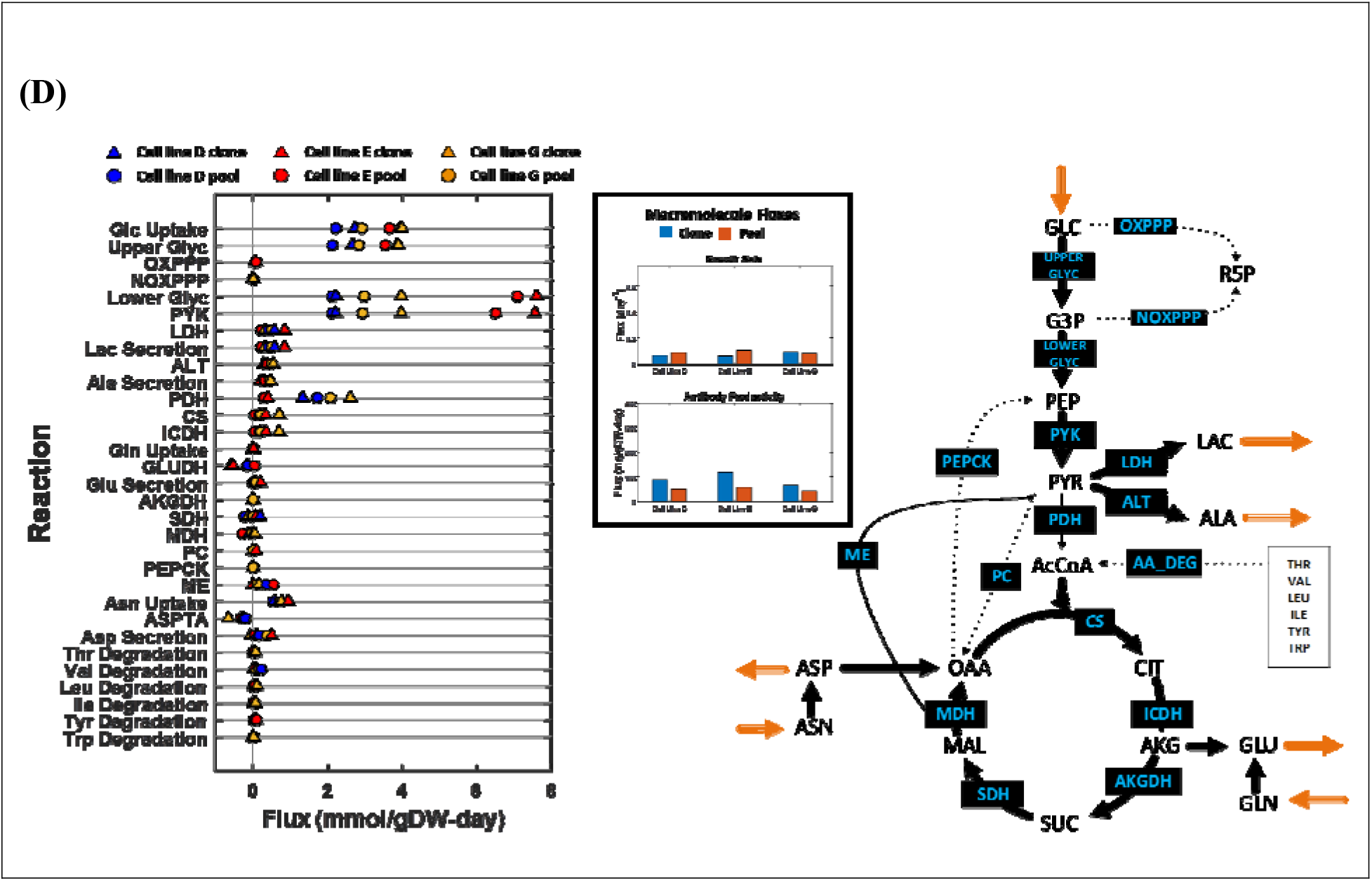
Flux maps showing the distribution of metabolic fluxes in the early exponential phase (A), lactate co-utilization phase (B), antibody production phase (C), and the late stationary phase (D). The arrows indicate the direction of metabolic flux (black arrows represent intracellular reactions, while orange arrows represent exchange reactions with the medium), metabolites are shown in white boxes, and reaction name abbreviations are shown in blue inside black boxes on the arrows. Fluxes (in mmol/gDW-day) are shown in graphs adjacent to the pathway maps. The orange and blue bars indicate the growth rate and antibody productivity for the pool and clone cultures, respectively.

The lactate co-utilization phase was marked by a reversal from lactate secretion to lactate consumption in all cell lines (Figure 5B). This was accompanied by a reduction in glucose uptake. Pools and clones showed similar metabolism during this phase with glucose being metabolized via the EMP pathway. The only exception here was the pool-G, which channeled all of the glucose through OXPPP. Since lactate is a major redox sink, the reversal from lactate secretion to lactate consumption leads to a reductive metabolic state, which drove flux into reduced metabolites such as glycerol and other fatty acid intermediates. Acetyl-CoA was synthesized from pyruvate via the PDH reaction, in all cases, in quantities sufficient to meet the demands for fatty acid biosynthesis. Consequently, the TCA cycle operated in the conventional direction and was further fueled by glutamine degradation. In addition, pyruvate carboxylase (PC) was active in all cases to replenish OAA and supplement aspartate biosynthesis over that produced by asparagine degradation.

The antibody production and late stationary phases exhibited a general slowdown of metabolism compared to the growth phases (Figure 5C and 5D) and accounted for much of the differences in process characteristics. Both phases showed reduced glucose uptake, reduced lactate secretion, and a significantly reduced uptake of all essential amino acids, due to nutrient depletion in the growth media. Metabolism in the ABP was streamlined to channel amino acids into antibody production in all cases in addition to biomass formation in clone-G. This contributed to the 1.7-fold higher growth rate in clone-G compared to pool-G, allowing a higher cell density in the clone culture. The cells minimally relied on metabolism to synthesize non-essential amino acids. Instead, aspartate, glutamate, and glycine secreted in the earlier phases of the process were taken up again by the cell and used for antibody production. Because the uptake of glutamate from the growth media was insufficient to meet the antibody synthesis demands, additional glutamate was synthesized from AKG via transaminase reactions, notably ASPTA, indicating that aspartate functioned as the primary amino group donor during the non-growth phases. The switch from aspartate, glutamate, alanine, and glycine production to consumption along with a near-zero degradation of essential amino acids suggests that amino acid uptake is a potential process bottleneck affecting antibody production during ABP.

### 3.5. Metabolic tasks are limited by amino acid uptake in clones

To better understand how amino acid availability impacted cell culture performance, we first computed the Pearson correlation coefficient (PCC) between all essential amino acid uptake rates and key metabolic tasks (growth rate and antibody specific productivity) in each cell line and phase (Supplementary Figure S2). Amino acid utilization did not always reflect the changes in growth rate and antibody production between the clone and pool cultures. In all cases, however, the uptake rates of essential amino acids (valine, leucine, isoleucine, phenylalanine, tyrosine, tryptophan, histidine, arginine, and methionine) strongly correlated with each other in all four phases (Supplementary Figure S2C). During the EXP and LCP, amino acids were abundantly available and therefore, did not limit the flux through the biomass formation reaction (Figure 6). Cell line E showed a positive correlation between growth rate and essential amino acid uptake in the early exponential growth phase, with clones showing a reduced amino acid utilization in this phase. Cell line D, however, showed a mixed trend in terms of essential amino acid utilization. Similar to cell line E, the clone cultures of cell line D grew slower and consumed smaller amounts of essential amino acids than the pool cultures in the early exponential phase. Because the reduction in growth rate was larger than the reduction in the uptake of essential amino acids, the clones could shunt more essential amino acids into their corresponding degradation pathways. Furthermore, degradation of essential amino acids generated acetyl-CoA as the end product, which reduced the reliance on reductive carboxylation for fatty acid biosynthesis (Figure 5A). In the early exponential and the lactate co-utilization phases for cell line G, growth rate negatively correlated with the uptake rate of essential amino acids, implying that faster growth did not necessitate increased amino acid uptake. This was because clone-G showed a 50% reduction in the uptake of valine, leucine, isoleucine, phenylalanine, and threonine without a significant change in the growth rate during this phase compared to pool-G. Thus, the uptake of these amino acids limited the flux through the biomass reaction in clone-G.

**Figure 6:**
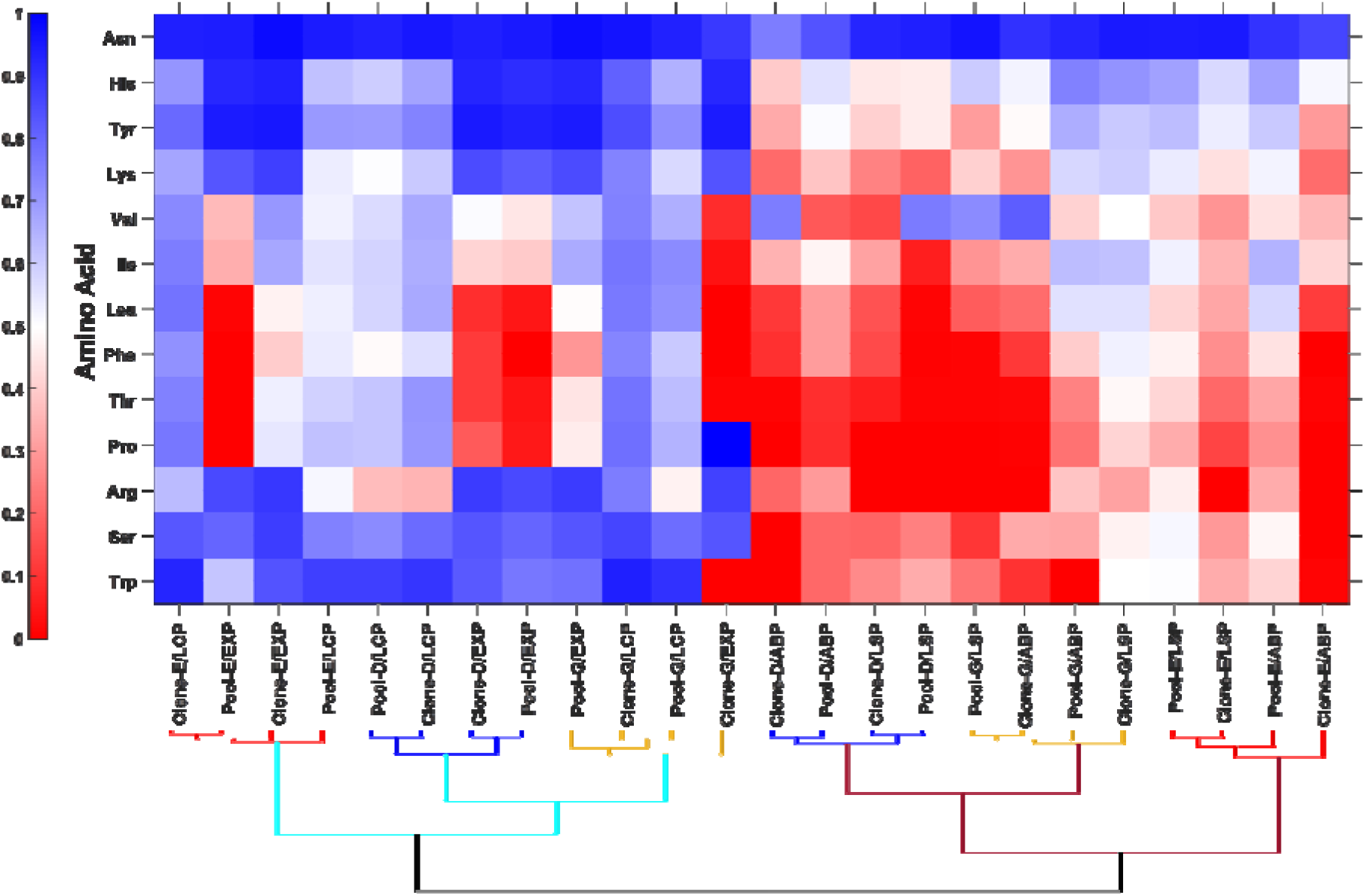
Heat map showing the fraction of consumed amino acids that were degraded via amino acid degradation pathways. On the heatmap, a blue color (corresponding to a value of 1) indicates an abundance of the amino acid over the minimum requirement for biosynthetic tasks in the specific condition and a red color (corresponding to a value of 0) indicates the consumed amino acid is completely channeled into biosynthetic tasks. In the dendrogram, cell line D i represented in blue, E in red, and G in orange. The cyan lines correspond to the growth phase models (early exponential and lactate co-utilization phases) whereas the maroon lines correspond to the non-growth phases (antibody production and the late stationary phases).

During the antibody production phase, the Pearson correlation coefficient (PCC) between antibody productivity and the uptake of key essential amino acids was between 0.4 and 0.6, implying that increased amino acid uptake only explained up to 36% of the observed increased antibody production in this phase. Uptake of most essential amino acids increased between 10% and 20%. The antibody productivity, however, increased by up to 3-fold in the clones. This was accomplished by diverting flux away from amino acid degradation and into antibody production in all clones. Flux sampling revealed that less than 10% of the consumed leucine, threonine, and proline were degraded in clone-D (down from 30%). On the other hand, cell line E shows a more efficient utilization of arginine, proline, phenylalanine, threonine, and tryptophan towards antibody production, leaving none for the degradation pathways. This suggests that the uptake of these amino acids limits the productivity of the antibody during the ABP. The same essential amino acids were limiting in clone-G. Contrary to the other essential amino acids, the uptake of valine increased by 5-fold in clone-D, by 1.6-fold in clone-E, and by 5-fold in clone-G compared to their respective pools during the ABP. This, consequently, significantly increased the flux through valine degradation in all cases as the uptake of valine vastly exceeded the demand for antibody production in all three clones. The same trend was seen in the LSP, but the magnitude of all fluxes was lower owing to reduced amino acid availability stemming from nutrient depletion. Overall, these findings show that by shifting fluxes away from amino acid degradation pathways, the high-producing clones achieved significantly higher antibody specific productivities than the corresponding pool cultures with only a modest increase in the uptake of these essential amino acids.

## 4. Discussion

This study presents a comprehensive exo-metabolomic, transcriptomic, and fluxomic characterization of three cell lines (including 3 stable pools and a high-producing clone derived from each pool) producing antibody to varying degrees. We found that high antibody titers achieved by high-producing clones depends on both specific productivity as well as cell density at the start of the antibody production phase. Because antibody production and cell growth are competing metabolic tasks, cell line optimization entails efficient channeling of resources towards desired metabolic tasks (cell growth and antibody production), minimization of byproducts, and controlled switching of metabolic priority from cell growth to antibody production at the right time. In this study, we found that rewiring nitrogen metabolism to take up and channel amino acids more efficiently into desired products was a hallmark of all high-producing clones. This included reduced nitrogen secretion via alanine, ammonia, aspartate, and glutamate, as well as parsimonious uptake of essential amino acids during the growth phases. Unlike previously reported findings (Templeton et al., 2013; Tharmalingam et al., 2018), we found that the final cell density did not positively correlate with high titer, implying that growth to high cell density, while desired, is not essential to a higher titer. Two of the three cell lines evaluated here had either the same or lower final cell density than their corresponding pool cultures. This suggests that antibody productivity had a stronger effect on final titer in cell lines D and E. Clone-G only showed a 50% improvement in antibody productivity compared to pool-G but resulted in a higher final titer than the pool culture by growing to a higher final cell density. These patterns are consistent with previous studies wherein clonal stability showed mixed effects of integral of viable cell density (IVCD) on titer (Dobson et al., 2020). Consistent with previously reported studies (Coulet et al., 2022; Mulukutla et al., 2015), we found multiple phases within the bioprocess, each associated with metabolic shifts. The cells prioritized growth between day 0 and day 6. However, they switched from lactate production to lactate consumption after day 3. This sub-phase within the growth phase has only been observed in batch and fed-batch cultures (Altamirano et al., 2006; Brunner et al., 2018), but not perfusion processes and has been attributed to lactate accumulation in the growth media (Torres et al., 2018). Elimination of lactate secretion is relevant to culture performance as lactate is a known inhibitor. This has been the focus of previous efforts to develop cell lines that reduce or eliminate lactate secretion (Jeon et al., 2011; Richelle and Lewis, 2017; Toussaint et al., 2016). Maximum antibody production was observed in the early stationary phase (antibody production phase) and was associated with a reuptake of previously secreted aspartate and glutamate. The late stationary phase exhibited a decline in metabolite uptake and antibody productivity, however, without a consistent loss of viability in all cell lines.

Monte-Carlo flux sampling provided valuable insights into phase-specific flux modes. We found that the overall carbon uptake declined over the course of the process (Figure 7A). However, a consistent quantity of carbon moles was allocated to unmeasured byproducts across conditions. Untargeted metabolomics (Gertsman and Barshop, 2018; Kao et al., 2021) in conjunction with stable-isotope tracing techniques (You et al., 2014) are valuable for identifying the currently unknown final products and their associated production pathways, although acetate, glycerol, sorbitol, fructose, citrate, fumarate, and malate have been previously identified as byproduct (Pereira et al., 2018). This information can be used alongside gene expression data to extract the most relevant context-specific models of metabolism. A key modality observed in all cases was the separation of cytosolic and mitochondrial metabolism. While cytosolic glycolysis via the EMP and pentose phosphate pathways was fueled by glucose, the TCA metabolism was fueled by glutaminolysis. Although a powerful tool to investigate mammalian metabolism given its ability to estimate all metabolic fluxes in a cell, flux sampling does not add any new constraints; therefore, it remains inferior to gold standard stable-isotope tracing techniques such as ^13^C- and ^15^N-metabolic flux analysis when applied to pathways that can resolved with those techniques. Previous tracer studies (Murphy et al., 2013) have already highlighted the need for a combination of positionally labeled tracers to sufficiently resolve split ratios such as the EMP/pentose phosphate pathway split in mammalian metabolism. The findings from this study highlight the necessity for stable-isotope tracers from multiple nutrients (e.g., glucose, glutamine, and asparagine) to elucidate different parts of metabolism. Furthermore, given the dynamic nature of metabolism in a bioprocess, accurate elucidation of metabolic shifts motivates the need for developing tracer data analysis paradigms that can handle both isotopic and metabolic instationary states.

**Figure 7A:**
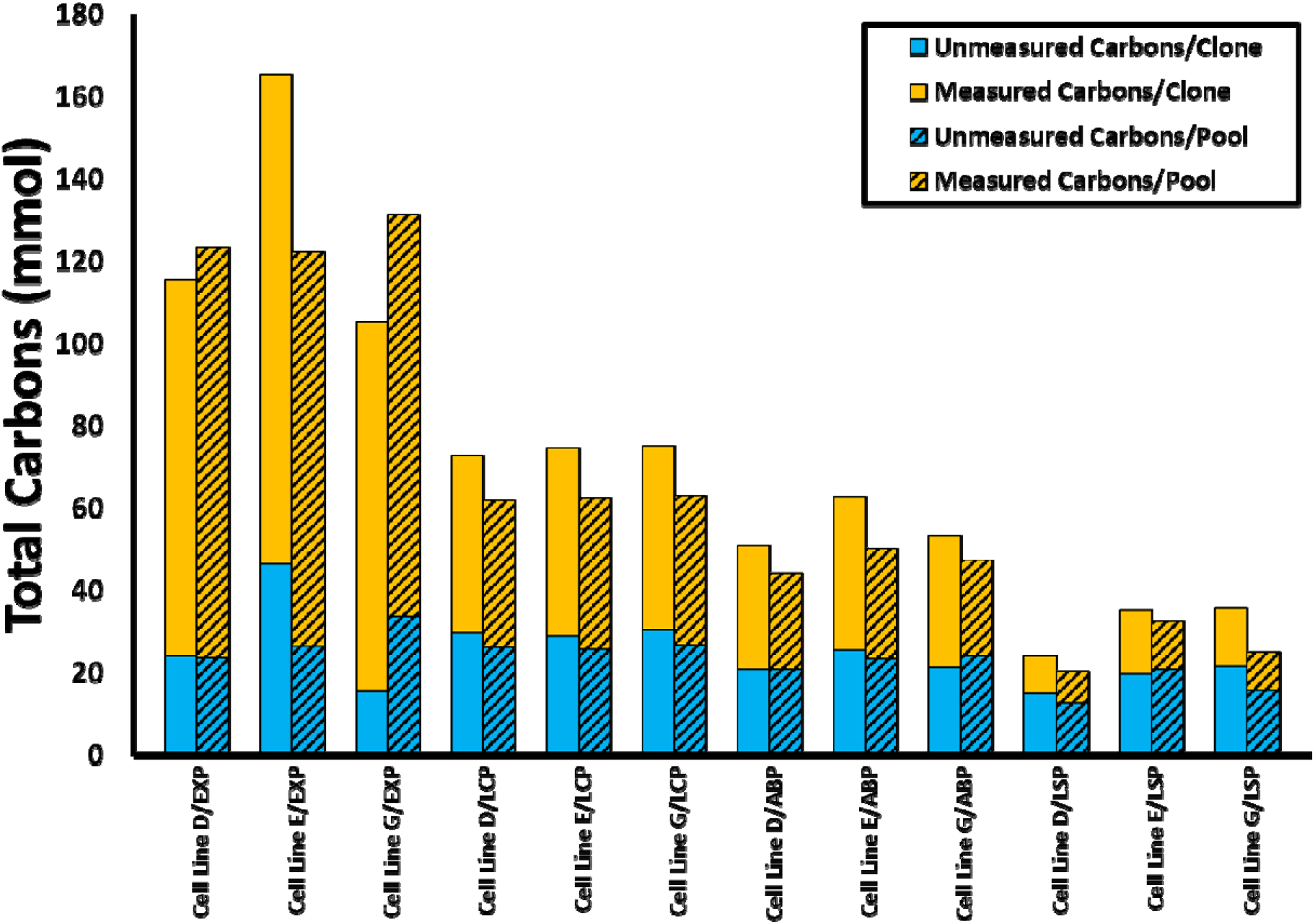
Carbon uptake and balance over the course of the bioprocess in each cell line and condition (with the stable pool and its best derivative clone plotted next to each other). The orange bars represent total carbons that are measured in end products including biomass, antibody, lactate, and other secreted amino acids. The blue bars represent carbons in unmeasured products.

Multi-omic analysis of the production process revealed that primary metabolic tasks are predominantly limited by the availability and uptake of amino acids. We found that the task efficiency (defined as the ratio of measured flux to the maximum flux predicted by the metabolic model) was higher for the clones of cell lines D and E in the production phase, indicating that these cells were already efficiently utilizing the consumed resources towards antibody production and that they are indeed limited by the uptake of limiting essential amino acids (Figure 6 and Figure 7B). The task efficiency in the growth phases was higher in the clone-G than pool-G, indicating that the observed increased final VCD was not merely a consequence of increased nutrient uptake, but due to efficient channeling of nutrients into biomass precursor pathways. Interestingly, the clones had a lower task efficiency in the antibody production phase because the nutrients were simultaneously used for cell growth and antibody production. The high task efficiency indicates that dominant metabolic tasks are limited by the availability of key metabolites. Because these limiting metabolites were not depleted in the media under these conditions, it can be argued that the uptake of these amino acids is a production bottleneck worthy of further investigation.

**Figure 7B:**
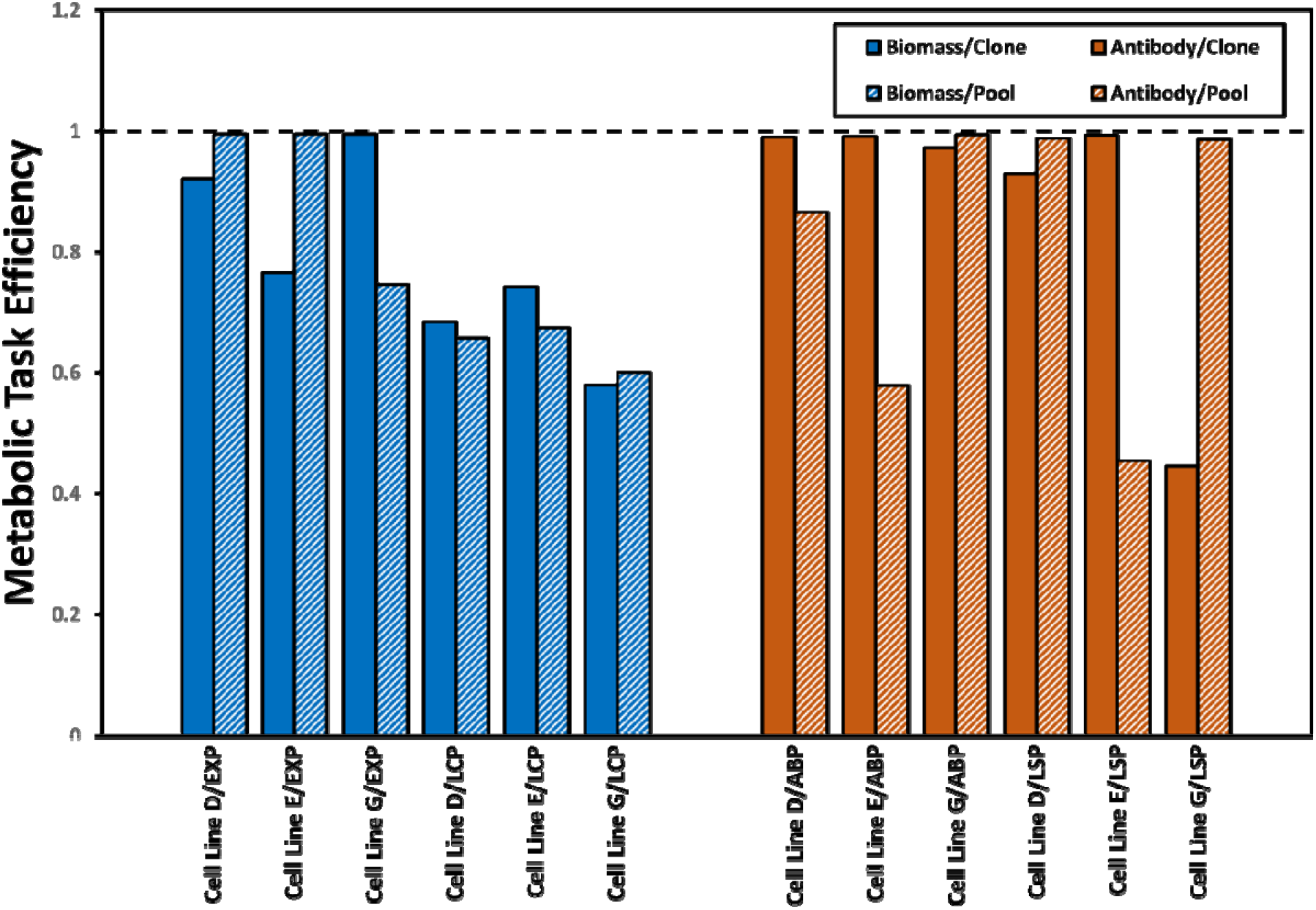
Efficiency of primary metabolic tasks defined as the ratio of measured flux to theoretical maximum flux predicted by the metabolic model. Biomass formation was considered to be the primary metabolic task in the EEP and LCP, whereas antibody production was considered the primary metabolic task in the ABP and LSP. An efficiency of 1 indicates that the task is only limited by the availability and/or uptake of precursor amino acids. A lower efficiency indicates resource usage towards competing byproducts.

A key question not addressed in this study is the factors and mechanisms underpinning cell state shifts in a bioprocess which led to the observed phase-specific metabolic properties. Our data indicate that the essential amino acids limiting cell growth and productivity are not depleted at any time during the process. Instead, asparagine and glutamine were found to be depleted. Interestingly, the availability of these amino acids did not affect the overall antibody productivity and titer. While beyond the scope of this dataset, concentration data from more frequent sampling can be analyzed to glean insights into bioreactor conditions that correlate with phase shifts. In addition, transcriptomics and protein-protein interaction studies (Samoudi et al., 2021) can elucidate the molecular mechanisms that facilitate condition sensing and subsequent cell-state shifts. These pathways can be attractive targets for cell-state modulation to trigger the shift from cell growth to antibody production when a sufficiently high VCD has been achieved in a bioreactor. Developing detailed cell-state models and integrating them with bioreactor models can be invaluable for holistic process modeling and optimization.

## Acknowledgements

This work was supported by funding generously provided by Amgen.

